# Simulation-informed low-current anodal tDCS accelerates early motor recovery after photochemically induced cortical stroke in rats

**DOI:** 10.64898/2026.08.04.742668

**Authors:** Saho Morishita, Satoshi Tanaka, Eikei Yamada, Akimasa Hirata, Tatsuro Kumada

**Author notes:** Correspondence should be addressed to: Dr. Tatsuro Kumada, Department of Occupational Therapy, Faculty of Health and Medical Sciences, Tokoha University, 1230 Miyakoda-cho, Hamakita-ku, Hamamatsu, Shizuoka, 431-2102, Japan.

## Abstract

Animal transcranial direct current stimulation (tDCS) studies typically use intensities exceeding clinical levels, and the off-line behavioral effects of weak electric fields in rodents remain unclear. We examined whether repeated anodal tDCS, calibrated by electric-field simulation to approximate human-equivalent weak fields, facilitates motor recovery after focal photothrombotic ischemic stroke (PIT) in rats. Simulation estimated that 50 μA produced a maximum field of ∼1.96 V/m in the targeted motor cortex, matching clinically relevant intensities. Under isoflurane anesthesia, rats received anodal tDCS at 50 μA, 250 μA, or 1 mA (5 min/day, 5 days/week, 2 weeks), or sham; motor recovery was assessed weekly by beam-walking for 4 weeks. A linear mixed-effects model revealed significant time, group, and time × group effects. The 50 μA group outperformed the PIT group at 1 week, and the 1 mA group at 2 weeks, with no differences thereafter. Low-current tDCS accelerates early post-stroke motor recovery, supporting weak-field neuromodulation.

## Introduction

Transcranial direct current stimulation (tDCS) modulates cortical excitability and induces neuroplastic changes by delivering weak electrical currents (1-2 mA) through scalp electrodes (Nitsche & Paulus, 2000; Stagg & Nitsche, 2011) . Meta-analyses have demonstrated significant beneficial effects on motor learning and functional recovery in patients with stroke, particularly when combined with rehabilitation training (Elsner et al., 2020; Kang et al., 2016). However, the precise neurophysiological mechanisms remain incompletely understood, necessitating the use of animal models for mechanistic investigations (Brunoni et al., 2011; Jackson et al., 2016).

A critical translational limitation has been the use of stimulation intensities in rodent studies far exceeding clinical levels (Liebetanz et al., 2006; T. Tanaka et al., 2013, 2020). Animal studies have employed current densities averaging 34.2 A/m^2^, over 85-fold higher than those used in human protocols (Brunoni et al., 2011) , raising serious concerns about translational validity (Jackson et al., 2016). Recently, Farahani et al. (2025) addressed this gap by demonstrating that repeated tDCS at clinically relevant field intensities (≈2 V/m) can enhance motor skill learning in awake rats when delivered concurrently with motor training, providing the first evidence that weak electric fields within the human therapeutic range produce measurable behavioral effects in rodents. However, this demonstration was confined to an online paradigm, in which stimulation was applied simultaneously with behavioral engagement. Whether clinically relevant weak-current tDCS can produce behavioral benefits when delivered offline, independently of concurrent behavioral activity, remains unknown. Equally, whether such human-equivalent weak fields can drive functional benefit in a clinically relevant disease context, rather than in the intact brain, remains untested. This distinction matters for both mechanistic and clinical reasons. Offline stimulation isolates activity-independent plasticity from learning-related processes, enabling cleaner investigation of neuromodulatory mechanisms (Kronberg et al., 2017; Turrigiano, 2017) . Clinically, it may benefit patients unable to participate in active training, including those with severe motor impairments or disorders of consciousness (Giacino et al., 2012).

To establish a translationally valid offline protocol, we performed computational electric-field simulation to calibrate current intensities in the rat brain to approximate the electric fields generated by standard clinical tDCS in humans. Based on this, we selected a primary low-current condition within the clinically relevant range, alongside higher intensities for comparison. Stimulation was administered under isoflurane anesthesia, with behavioral outcomes assessed separately, in a photothrombotic ischemic stroke (PIT) rat model targeting the motor cortex. Our aims were to determine whether simulation-calibrated offline tDCS facilitates post-stroke motor recovery and to provide a robust platform for investigating the cellular mechanisms of weak-field neuromodulation.

## Methods

### Animals and ethics, and photothrombotic ischemic stroke (PIT)

Adult Sprague–Dawley rats (8–10 weeks old, 250–320 g) were used in this study. All experimental procedures were approved by the Animal Care and Use Committee of Tokoha University. Focal motor cortical infarction was induced by the photothrombotic method, and infarct formation was verified by postoperative motor deficits and TTC staining, as previously described (Morishita et al., 2020).

### Electric-field simulation and current intensity selection

To establish a stimulation protocol with translational relevance to human studies, a computational electric-field simulation was performed to estimate the stimulation current required to generate human-equivalent electric fields in the rat brain. A computational rat model (body weight: 265 g) was constructed from CT images with a spatial resolution of 0.25 mm and segmented into six tissue types: skin, muscle, fat, cortical bone, brain (grey matter), and eye (Figure 1A-i). Each tissue was modeled as a linear isotropic conductor, and tissue conductivities were assigned according to experimentally reported values for the corresponding human tissues (Nishimoto et al., 2024). The conductivity values used in the simulation are listed in Table 1. Conductivities of the electrode materials were assigned according to Laakso et al. (2016). A 12-mm-diameter metal electrode (anode) was positioned over the left primary motor cortex (M1), while a 3 × 3 cm rubber electrode (cathode) was placed on the ventral thorax (Figure 1A-ii). The induced electric field was calculated using the scalar-potential finite-difference (SPFD) method (Dawson & Stuchly, 1998; Laakso et al., 2016) . Under the quasi-static approximation, the scalar potential φ was obtained by solving

**Table 1.**
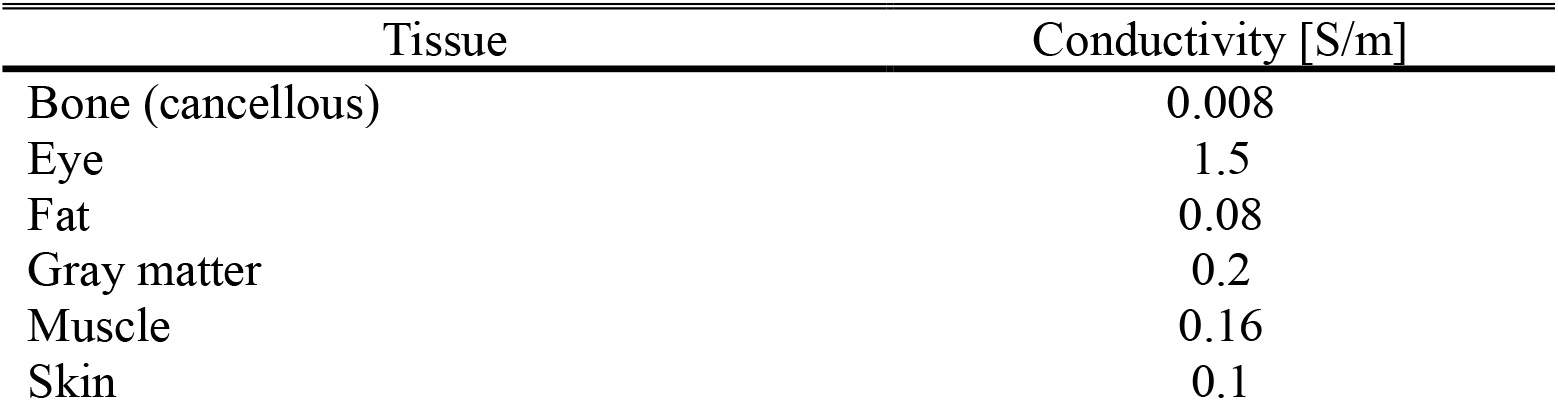

**Figure 1.**
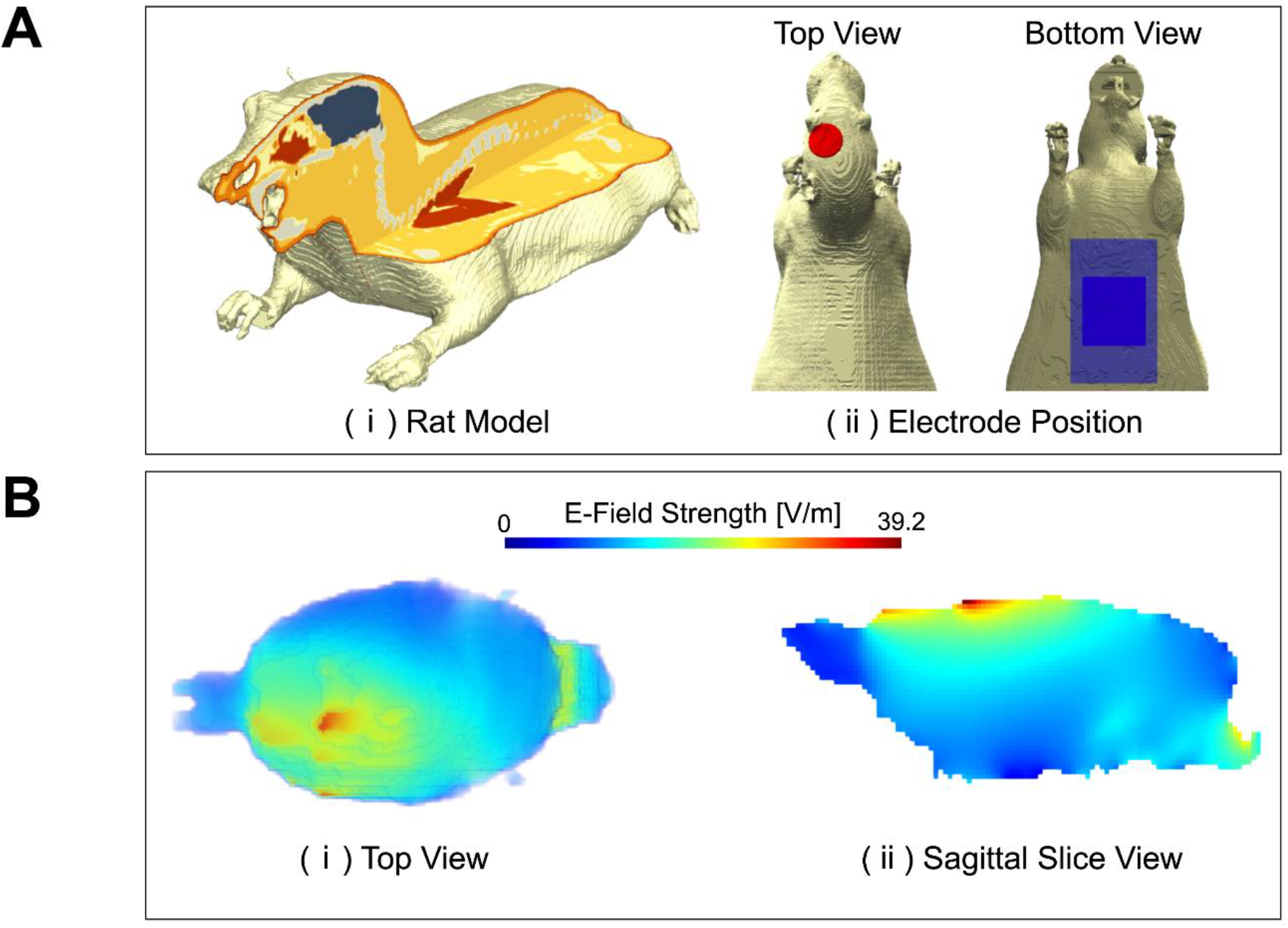
Computational model and simulated electric field distribution for low-current tDCS. (A) Rat model and electrode placement. (i) Segmented rat model showing the six tissue types used in the simulation. (ii) Electrode positions: the anode (red) over the left primary motor cortex (M1) and the cathode (blue) over the ventral thorax. (B) Simulated electric-field magnitude in the grey matter at an injection current of 1 mA. (i) Top view of the field distribution. (ii) Sagittal slice through the left M1. The maximum field in the left M1 was 39.2 V/m. Color bar, electric-field magnitude (V/m).

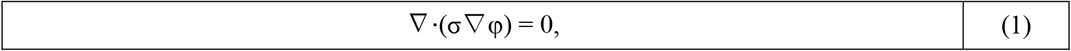

where σ denotes the electrical conductivity. The scalar potential was discretized on a voxel grid, and electrical conductance between neighboring nodes was determined from the local conductivity distribution. Application of Kirchhoff’s current law at each node yielded a sparse system of linear equations, which was solved using a multigrid algorithm with successive over-relaxation (SOR). Six multigrid levels were used, and iterations continued until the relative residual fell below 10^−6^ (Laakso & Hirata, 2012). The electric field was then calculated from the potential difference between adjacent nodes divided by the inter-node distance. The simulation predicted a maximum electric field of 39.2 V/m within the targeted left M1 region for a current injection of 1 mA (Figure 1B). Assuming a linear relationship between injected current and induced electric field, stimulation currents of 50 μA and 250 μA were estimated to generate maximum electric fields of approximately 1.96 V/m and 9.79 V/m, respectively. Because the electric field estimated for 50 μA was comparable to cortical electric-field magnitudes reported in clinically relevant human tDCS dosimetry studies (Farahani et al., 2025), 50 μA was selected as the primary low-current stimulation condition. Higher-current conditions (250 μA and 1 mA) were included to evaluate intensity-dependent effects.

### tDCS procedure and behavioral assessment

Rats were assigned to the following groups: sham, PIT, PIT + 50 μA tDCS, PIT + 250 μA tDCS, and PIT + 1 mA tDCS. During deep anesthesia with isoflurane, the stimulating electrode was placed on the scalp over the ipsilateral cerebral cortex, and a reference rubber electrode (3 × 3 cm) was placed on the shaved chest using adhesive conductive Ten20 paste (Weaver and Company, Aurora, CO, USA). Anodal tDCS was delivered continuously for 5 min using a constant current stimulator (DC-Stimulator Plus, NeuroConn, Germany) at 50 μA, 250 μA, or 1 mA. Stimulation was administered five days per week for two weeks.

Motor coordination was evaluated using the beam-walking test on an elevated wooden beam, with performance rated on a 0–5 ordinal scale based on foot slip frequency, as previously described (Morishita et al., 2020). Testing was performed at baseline before surgery (-1W), immediately after PIT (0 W), and weekly for 4 weeks.

### Statistical analysis

Beam-walking scores, rated on a 0–5 ordinal scale, are presented as median with interquartile range. For paired within-subject comparisons between baseline (-1W) and the time point immediately after PIT (0W), the Wilcoxon matched-pairs signed-rank test was used. To assess the time course of motor recovery across stimulation groups, beam-walking scores from 1 to 4 weeks after PIT were analyzed using a mixed-effects model (restricted maximum likelihood estimation), with time as a within-subject factor and group as a between-subject factor; the time × group interaction was evaluated to test whether recovery trajectories differed among groups. At each post-PIT time point, between-group comparisons against the PIT-only group were performed using the Kruskal–Wallis test followed by Dunn’s multiple comparisons test, with adjusted *p* values reported. Animals were allocated to stimulation groups in a pseudorandom order. Behavioral scoring was performed by a single trained investigator using the predefined ordinal criteria. All statistical analyses were performed using GraphPad Prism version 9.5.1 (GraphPad Software, San Diego, CA, USA), and statistical significance was set at *p* < 0.05.

## Results

### Validation of the PIT stroke model

To investigate the effects of low-current tDCS on motor recovery, we used a rat model of focal cerebral infarction in the motor cortex induced by photothrombotic ischemic stroke (PIT) (Figure 2A). Consistent with our previous report (Morishita et al., 2020), TTC staining of coronal brain sections showed that the infarct was primarily localized to a narrow medial parietal cortical region corresponding to the limb area of the motor cortex, whereas the subcortical regions appeared largely preserved (Figure 2B). Behaviorally, all animals included in the analysis exhibited motor impairments after PIT. Beam-walking scores at 0 W were significantly worse than baseline scores at -1 W (Wilcoxon matched-pairs signed-rank test, *p* < 0.0001), confirming the successful induction of the PIT model.

**Figure 2.**
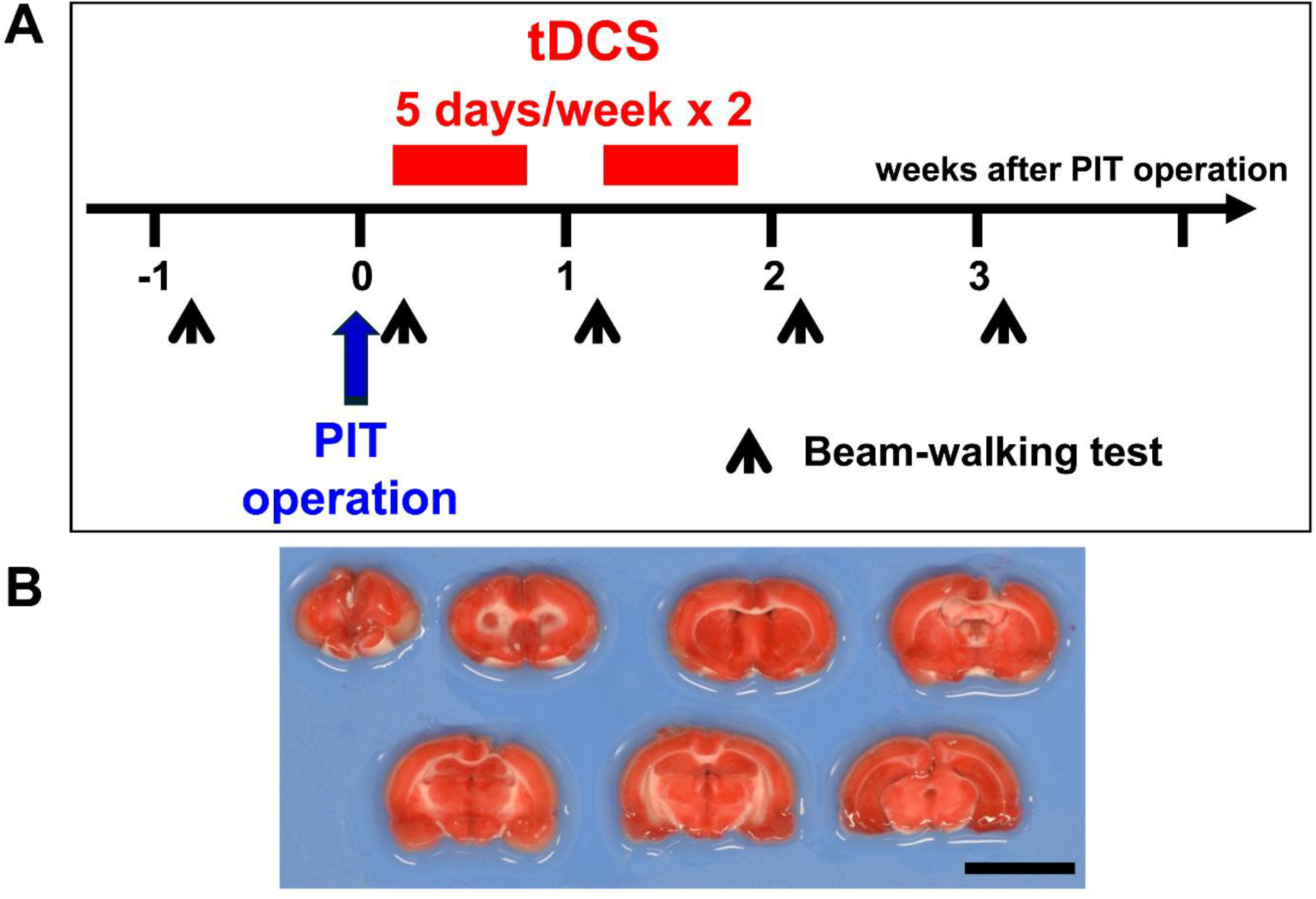
Experimental design and validation of the photothrombotic ischemic (PIT) stroke model. (A) Schematic timeline of the study. Following PIT induction (0W), low-current transcranial direct current stimulation (tDCS) was applied 5 days/week for 2 weeks. Black downward arrows indicate the timing of the beam-walking test. (B) Representative TTC-stained coronal brain sections at 4 weeks post-PIT. Viable tissue appears red, while infarcted areas appear white.

### Low-current tDCS accelerates early motor recovery after PIT

Motor recovery was longitudinally assessed for 4 weeks using a beam-walking test (Figure 3A). Mixed-effects analysis revealed significant main effects of time (F (2.88, 23.04) = 65.67, *p* < 0.0001), group (F (2.46, 19.71) = 4.19, *p* = 0.0241), and time × group interaction (F (4.70, 26.29) = 5.28, *p* = 0.0020), indicating that the recovery time course differed among the experimental groups. Across groups, beam-walking performance showed a biphasic pattern, with an improvement up to 2 weeks after PIT, a transient decline at 3 weeks, and partial recovery at 4 weeks.

**Figure 3.**
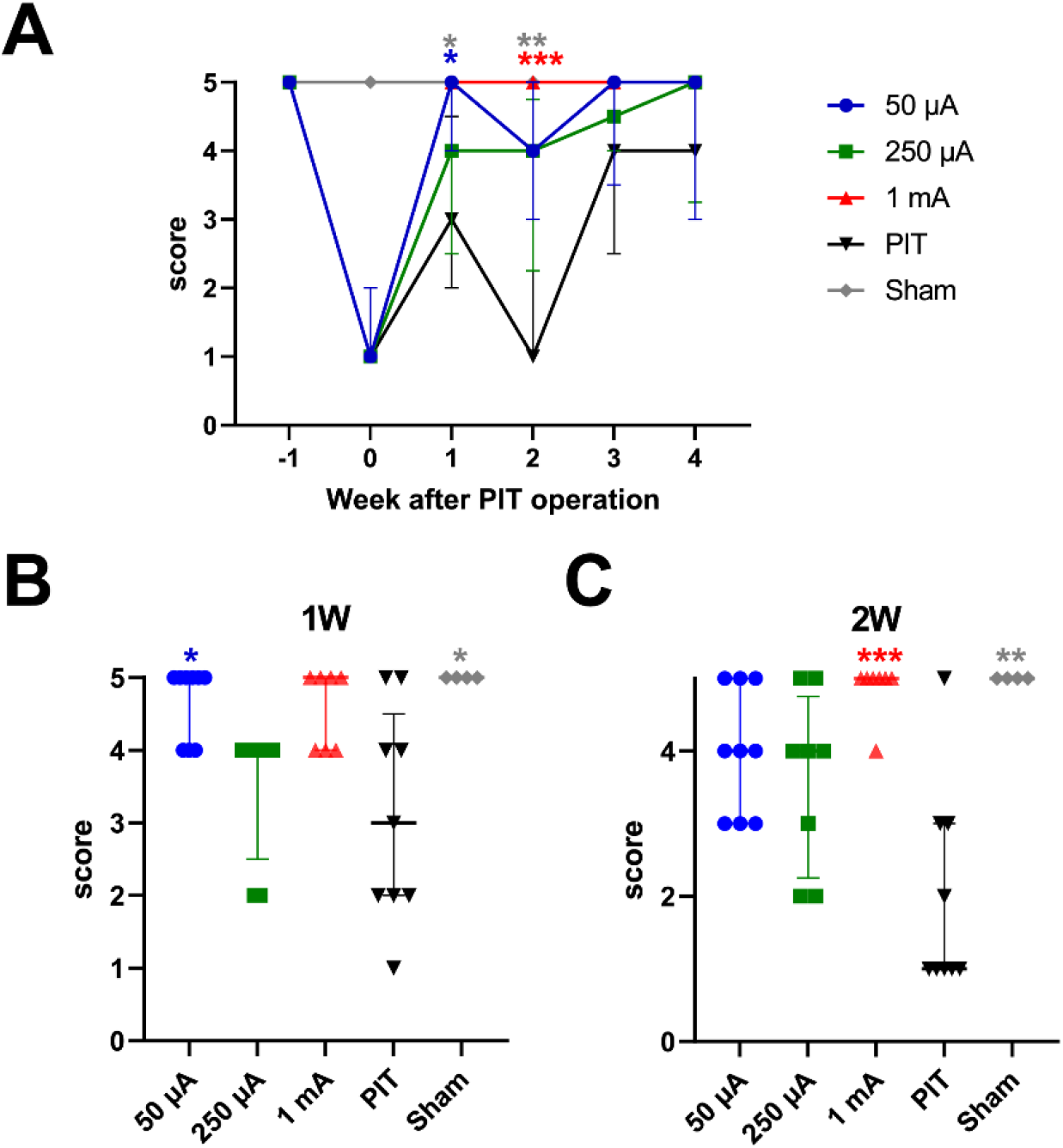
Low-current tDCS accelerates early functional motor recovery. (A) Time course of beam-walking test scores from pre-operation to 4 weeks post-PIT. Data are presented as median with interquartile range. A mixed-effects model revealed a significant time × intervention interaction (p = 0.0020). (B, C) Scatter plots showing individual beam-walking scores at 1 week (B) and 2 weeks (C) post-PIT. Statistical significance was determined using the Kruskal-Wallis test followed by Dunn’s multiple comparisons test (*p < 0.05, **p < 0.01, *** p < 0.001 vs. PIT).

At 1 week after PIT, the 50 μA tDCS group showed significantly higher beam-walking scores than the PIT group (Dunn’s multiple comparisons test, adjusted *p* = 0.0335; Figure 3B). In addition, the sham group showed significantly higher scores than the PIT group at this time point. At 2 weeks after PIT, the 1 mA tDCS group showed significantly higher scores than the PIT group (adjusted *p* = 0.0006; Figure 3C), and the sham group remained significantly different from the PIT group. No significant improvement was observed in the 250 μA group at either 1 week (adjusted *p* > 0.9999) or 2 weeks (adjusted *p* = 0.2844) compared to the PIT group. In contrast, no significant differences were detected between the tDCS-treated groups and the PIT group at 3 or 4 weeks after PIT.

Together, these findings indicate that low-current tDCS facilitates motor recovery predominantly during the early post-stroke phase.

## Discussion

The present study demonstrates that repeated low-current transcranial direct current stimulation (tDCS) can facilitate early motor recovery after focal cortical infarction, even when delivered offline under isoflurane. In the PIT model targeting the motor cortex, longitudinal analysis of beam-walking performance revealed significant effects of time, group, and time × group interactions. Time-point analyses further showed that the 50 μA group exhibited significantly higher beam-walking scores than the PIT group at 1 week, whereas the 1 mA group showed significantly higher scores at 2 weeks than the PIT group. In contrast, no significant differences were detected between the tDCS-treated groups and the PIT group at 3 or 4 weeks. Together, these findings suggest that the principal effect of low-current tDCS in this model is not to raise the final level of motor performance but rather to accelerate recovery during the early post-stroke period.

These findings are important in light of the recent report by Farahani et al. (2025), which showed that repeated tDCS at a clinically relevant field intensity can enhance motor learning in rats when stimulation is delivered concurrently with behavioral training. The overall intent of that study and ours is highly similar: both address the long-standing gap between the weak electric fields used in human tDCS and the substantially stronger stimulation commonly used in rodent studies. In this regard, the present results support and extend that emerging framework by showing that weak-current stimulation can also influence behavior in a distinct experimental context, namely post-stroke motor recovery. At the same time, the two studies differ in an important way. Farahani et al. focused on motor skill learning in non-lesioned awake behaving rats with concurrent stimulation, whereas the present study examined recovery after focal cortical infarction (a clinically relevant disease model) using repeated off-line stimulation under anesthesia. Our results therefore broaden the conditions under which weak-current tDCS may exert biologically meaningful effects. To our knowledge, this is the first demonstration that human-equivalent weak electric fields are sufficient to produce a measurable functional benefit in a post-stroke brain, extending the human-calibrated dosimetric paradigm validated by Farahani et al. (2025) from intact to injured cortex. Critically, the 50 μA condition used here was not chosen merely as a low absolute current, but as part of a simulation-informed attempt to approximate human-equivalent weak-current dosimetry; this current corresponded to a maximum field of approximately 1.96 V/m in the targeted left M1, comparable to the ≈2 V/m intensity reported by Farahani et al. (2025). Thus, the behavioral benefit observed at 50 μA provides experimental support for the biological relevance of microampere-level stimulation within a human-equivalent dosimetric framework.

The present data also suggests that weak-current tDCS may act by modulating recovery-related plasticity, rather than by directly driving neuronal activity. In lesioned motor circuits, subtle shifts in excitability, synaptic gain, or network states may be sufficient to influence the efficiency of spontaneous recovery mechanisms, consistent with the idea that weak electric fields may bias ongoing reorganization in peri-infarct networks without requiring overt, suprathreshold activation (Kronberg et al., 2017). Consistently, previous studies in rodent models have shown that microampere-range stimulation can influence neuroinflammation and functional recovery, supporting the biological relevance of weak-current neuromodulation (Huang et al., 2021; Peruzzotti-Jametti et al., 2013; Walter et al., 2022).

Notably, the effects were phase-specific rather than uniformly monotonic with current intensity. The earliest significant difference was observed in the 50 μA group at 1 week, whereas the strongest difference at 2 weeks was observed in the 1 mA group. These findings suggest that the effective stimulation range may depend on recovery stage, tissue state, or ongoing reorganization within peri-infarct networks, raising the possibility that the biologically effective range of tDCS after stroke may be narrower and more state-dependent than has often been assumed. From a translational perspective, this off-line protocol may be relevant for clinical situations in which active training is difficult, such as in patients with severe motor deficits. Importantly, awake-animal tDCS paradigms can be confounded by attention, motivation, and arousal state (Nitsche et al., 2003; Stagg & Nitsche, 2011), which our isoflurane-anesthetized off-line protocol inherently minimizes; this design also parallels evidence that stimulation delivered during rest or sleep can prime motor learning and memory consolidation (Marshall et al., 2004; Reis et al., 2008). Clinical tDCS over the motor cortex has been associated with improvements across multiple motor functions in humans (Kang et al., 2016; Reis et al., 2008; Stagg & Nitsche, 2011; S. Tanaka & Watanabe, 2009; S. Tanaka et al., 2011), supporting the translational relevance of the present off-line approach.

Several limitations should be acknowledged. First, motor outcome was assessed solely by the beam-walking test, and whether similar effects extend to other motor or cognitive domains remains to be determined. Second, the present study did not directly measure the neurophysiological, synaptic, or molecular mechanisms underlying the behavioral benefit of low-current tDCS. Third, because stimulation was delivered under isoflurane anesthesia, potential interactions between anesthesia and tDCS on the observed effects cannot be excluded. Fourth, the computational model used a generic rat anatomy and did not account for individual anatomical variation; subject-specific dosimetry could further refine the precision of weak-field neuromodulation in future studies. Fifth, because animal husbandry and behavioral scoring were performed by the same investigator, formal blinding to group allocation was not feasible; future studies with independent assessors blinded to group allocation will be needed to confirm the robustness of these findings.

## Conclusions

The present study shows that repeated offline anodal tDCS, calibrated via electric-field simulation to approximate clinically relevant field intensities, facilitates early motor recovery after focal cortical infarction in rats. Notably, the behavioral effect was observed at 50 μA, a condition estimated to generate a maximum electric field of approximately 1.96 V/m in the target motor cortex, comparable to the clinically relevant intensity recently validated in an online paradigm by Farahani et al. (2025). This finding extends this framework from intact to injured cortex by showing that behaviorally meaningful effects can be obtained in a clinically relevant focal stroke model even without concurrent behavioral engagement. Together, these results provide a translationally valid, simulation-informed offline platform for investigating the cellular and molecular mechanisms through which weak-current tDCS modulates post-stroke neural plasticity.

## Acknowledgements

The authors thank Dr. Jose Gomez-Tames for valuable advice on the computational electric-field simulation. The authors also thank all members of the laboratory for their technical assistance and helpful discussions.

During the preparation of this manuscript, the authors used Notion AI (Notion Labs, Inc.) and ChatGPT (OpenAI) for iterative manuscript revision, English language refinement, reference formatting, and editorial assistance based on co-author feedback. The authors also used Paperpal (Cactus Communications) for scientific language editing and proofreading. After using these tools, the authors reviewed and edited the content as needed and take full responsibility for the content of the publication.

## Author Contributions

**Saho Morishita**: Conceptualization, Methodology, Validation, Formal analysis, Investigation, Data curation, Writing – original draft, Writing – review & editing, Visualization. **Satoshi Tanaka**: Conceptualization, Methodology, Resources, Writing – original draft, Writing – review & editing, Supervision. **Eikei Yamada**: Methodology, Software, Validation, Writing – review & editing, Visualization. **Akimasa Hirata**: Methodology, Software, Resources, Writing – review & editing, Supervision. **Tatsuro Kumada**: Conceptualization, Methodology, Validation, Formal analysis, Investigation, Resources, Data curation, Writing – original draft, Writing – review & editing, Visualization, Supervision, Project administration, Funding acquisition. All authors reviewed and approved the final manuscript.

## Statement and Declarations

### Ethical considerations

All experimental procedures involving animals were approved by the Animal Care and Use Committee of Tokoha University and were performed in accordance with the institutional guidelines for the care and use of laboratory animals and the ARRIVE guidelines.

### Consent to participate

Not applicable.

### Consent for publication

Not applicable.

### Data availability statement

The data that support the findings of this study are available from the corresponding author upon reasonable request. The computational simulation code used in this study is available from the corresponding author upon reasonable request.

